# Motivating Effects of Negative-hedonic Valence Encoded in Engrams

**DOI:** 10.1101/2025.02.21.639600

**Authors:** Hermina Nedelescu, Elias Meamari, Nami Rajaei, Alexus Grey, Ryan Bullard, Nathan O’Connor, Nobuyoshi Suto, Friedbert Weiss

**Affiliations:** Department of Neuroscience, The Scripps Research Institute, 10550 North Torrey Pines Road, La Jolla, CA 92037; MBF Bioscience, 185 Allen Brook Ln 101, Williston, VT 05495; Department of Molecular Pharmacology and Experimental Therapeutics, Mayo Clinic, 200 First St. SW, Rochester, MN 55905

**Keywords:** Engrams, positive reinforcement, negative reinforcement, reward dysregulation, learning

## Abstract

Engrams are neuronal alterations that encode associations between environmental contexts and subjectively rewarding or aversive experiences within sparsely activated neuronal assemblies that regulate behavioral responses. How positive- or negative-hedonic states are represented in brain neurocircuits is a fundamental question relevant for understanding the processing of emotionally meaningful stimuli that drive appropriate or maladaptive behavior, respectively. It is well-known that animals avoid noxious stimuli and experiences. Little is known, however, how the conditioning of environmental or contextual stimuli to behavior that leads to amelioration of dysphoric states establishes powerful associations leading to compulsive maladaptive behavior. Here we have studied engrams that encode the conditioned effects of alcohol-related stimuli associated with the reversal of the dysphoric withdrawal state in alcohol dependent rats and document the recruitment of engrams in the paraventricular nucleus of the thalamus (PVT), the central nucleus of the amygdala (CeA), and the Dorsal Striatum (DS). The findings suggest that the encoding of associations between reversal of negative hedonic states and environmental contexts in these engrams may serve as a neural mechanism for compulsive alcohol seeking and vulnerability to relapse associated with dysregulation of reward to a pathological allostatic level.

## INTRODUCTION

The conditioning of environmental stimuli or contexts with rewarding or aversive events is an important process that becomes encoded in neurocircuits to drive behavior (Kim et al. 2016; Josselyn and Tonegawa 2020; Tye 2018; George and Hope 2017). Animals perceive stimuli as rewarding or aversive, they can also learn through associative processes that avoiding or removing negative stimuli ameliorates unpleasant experiences. In drug dependent subjects, for example, learned associations between contextual stimuli and the drug include associations linked to the reversal of the adverse withdrawal state by drug use. The ability to process meaningful stimuli related to this negative reinforcement learning is a vital neural function that is essential for maintaining stability, well-being and survival. How stimuli that drive behavior are represented in neurocircuits is, therefore, a fundamental question.

Exposure to stimuli elicits sparse patterns of neuronal activation known as neuronal ensembles, and the repetitive exposure to stimuli that encode long-lasting learned associations are known as engrams (Cruz et al. 2015; Cruz et al. 2013; Josselyn and Tonegawa 2020; Eichenbaum 2016; Hebb 2005; Cai et al. 2016). Since learned associations are a major factor in the chronically relapsing nature of compulsive alcohol seeking and use, the withdrawal-related learning (WDL) paradigm was employed to identity brain engrams that support the acquisition of this negative reinforcement learning and the development of alcohol dependence (Kozanian et al. 2022). In the WDL paradigm, environmental stimuli conditioned to learning about the reversal of aversive withdrawal states – negative reinforcing – of a substance (e.g., ethanol) during withdrawal acquire conditioned incentive value of their own and represent an independent major factor in substance craving and relapse. The WDL model provides an effective tool to advance current understanding of how behaviorally relevant learning experiences are established in functional neurocircuits containing engrams.

The formation of engrams that encode long-lasting learned associations which then mediate appetitively motivated behavior including drug-seeking responses are thought to be the basic unit of acquired learning and memory (Alvarez and Eichenbaum 2002; Cruz et al. 2015; Cruz et al. 2013; Pettit et al. 2022; Laque et al. 2019; Nedelescu et al. 2022; Josselyn and Tonegawa 2020; Suto et al. 2016). However, the neuronal substrates that specifically mediate the motivating effects of stimuli associated with the reversal of negative hedonic states such as dysphoria, anxiety, and sensitivity to stress after excessive or long-term substance use remain to be understood. The relevance of this understanding is illustrated by findings of a parallel behavioral study in which alcohol seeking induced by contextual stimuli associated with the reversal of adverse withdrawal states by alcohol (withdrawal-related learning [WDL]) was qualitatively and quantitatively different from that in rats without a dependence and WDL history (Kozanian et al. 2022). Specifically, alcohol seeking in rats with WDL experience was overall stronger and, more importantly, compulsive in nature (i.e., resistant to punishment and increased effort requirements) (American Psychiatric Association 2022), whereas alcohol seeking in nondependent rats with a “social drinking” history was not. Therefore, using rat reinstatement models we sought to (a) identify engrams that encode associations between environmental stimuli linked to alcohol availability and consumption during withdrawal episodes following the development of alcohol dependence (negative reinforcement) versus alcohol availability in the nondependent state (positive reinforcement), and (b) establish whether alcohol seeking in rats with WDL experience recruits different engrams than in rats without this experience as well as the nature of these differences.

## MATERIALS AND METHODS

### Animals

All procedures for the care of animals were in adherence to the NIH *Guide for the Care and Use of Laboratory Animals*, and approved by the Institutional Animal Care and Use Committee of Scripps Research. All animals were adult male Wistar rats (Charles River, Wilmington, MA) weighing approximately 450 grams.

Brains were obtained from rats trained and tested under the conditions elaborated below (Kozanian et al. 2022).

### Behavioral training

The withdrawal-related learning (WDL) behavioral paradigm and alcohol seeking under conditions of motivational and environmental challenges were previously described (Gonzalez-Cuevas et al. 2018; Kozanian et al. 2022). Briefly, WDL establishes the effects of contextual stimuli conditioned to alcohol reinforcement during withdrawal on reinstatement involving motivational and environmental challenges (**Figure 1**). Rats were divided into four groups (DEP-WDL; NDEP-WDL; DEP-NWDL; NDEP-NWDL), two of which were subjected to alcohol dependence (DEP-WDL and DEP-NWDL) induction while the other two were non-dependent (NDEP-WDL and NDEP-NWDL) (Vendruscolo and Roberts 2014). In phase I, all rats were trained to self-administer alcohol for three weeks. Following self-administration, in phase II, rats were subjected to alcohol vapor inhalation (DEP) or remained nondependent (NDEP). After three weeks, DEP-WDL or NDEP-WDL control rats were transiently removed from the vapor (or control) chambers and after eight hours of withdrawal given the opportunity to operantly self-administer alcohol for 30-minutes in the presence of an environmental context consisting of an auditory (tone) and an olfactory (anise scent) stimulus. Note that in the NDEP-WDL condition “WDL” refers procedurally to the same contextual stimulus exposure as in the DEP-WDL condition but does not imply withdrawal-related learning because these animals remained nondependent. Two additional control/comparison conditions were included. First, a “dependent no withdrawal-related learning” group (DEP-NWDL) that served the purpose of providing a comparison for the effects of a dependence history alone without withdrawal learning history. Following dependence induction, rats in this group were removed from the vapor chambers for one week to complete alcohol withdrawal in their home cages and then, in the post-dependent state, given the opportunity to operantly self-administer alcohol in the presence of same contextual stimuli as in the DEP-WDL condition. In parallel, an additional control group not exposed to alcohol vapor and thus remaining nondependent (NDEP-NWDL) was tested to compare the effects of alcohol-contextual stimulus associations acquired in the post-dependent (DEP-NWDL) state to that in rats trained and tested along the same timeline but remaining nondependent. After removal from the vapor or control chamber, rats in all groups were allowed to recover for 1 week before re-exposure to the self-administration chambers under extinction conditions and measurement of responses at the previously active lever following presentation of the respective alcohol associated stimulus context (described in (Kozanian et al. 2022)

**Figure 1.**
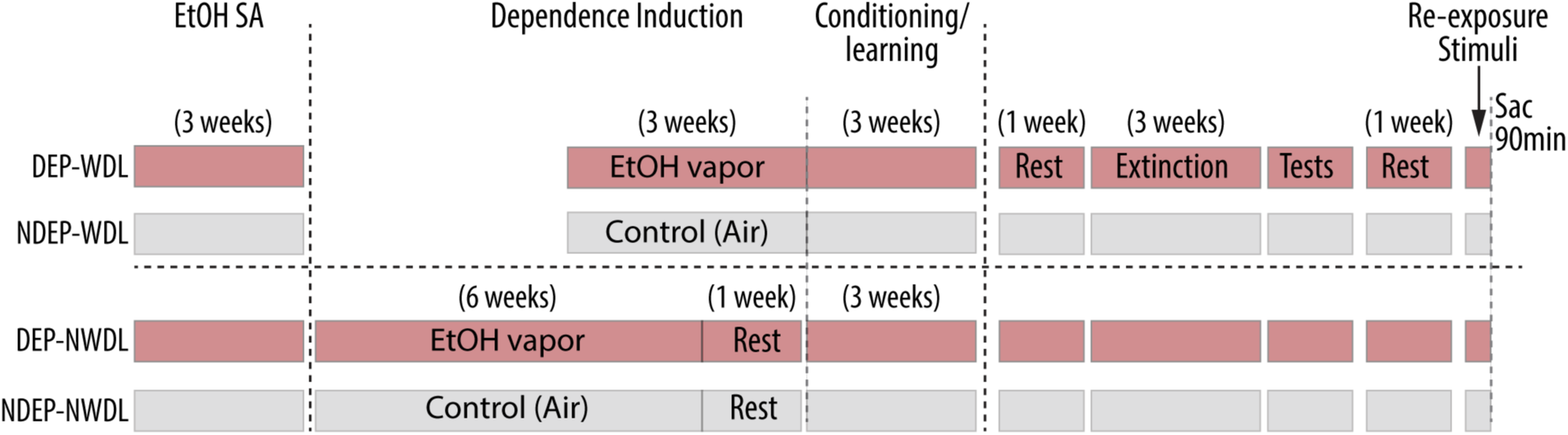
Illustration of the design including the withdrawal-related learning (WDL) procedure. Rats were divided into four groups (DEP-WDL; NDEP-WDL; DEP-NWDL; NDEP-NWDL) two of which were subjected to alcohol dependence (DEP-WDL and DEP-NWDL) induction while the other two remained non-dependent (NDEP-WDL and NDEP-NWDL). For explanations of the four test conditions and detailed description of training and test procedures, see Methods: Behavioral Training.

### Preparation of tissue for microcopy

Whole brains from our previous behavioral experiment (Kozanian et al. 2022) were used to analyze neuronal activation patterns in the present study. Briefly, tissue preparation was as follows: all rats were sacrificed with CO_2_ and transcardially perfused (4% paraformaldehyde in 0.1-mM sodium tetraborate) 90-minutes after re-exposure to contextual stimuli (without alcohol availability) in order to visualize Fos protein expression of activated neurons within engrams. Then, brains were harvested and placed in 30% sucrose before cutting 40-50 µm thick tissue sections using a Leica microtome. In this study, whole brains were sectioned at the same thickness and all tissue sections for this study were collected together into appropriate wells in a 24-well plate for immunostaining. Immunolabeling was achieved with an anti-cfos antibody (Cell Signaling, #2250 cfos (9F6) rabbit mAB, 1:5000). First, tissue sections were incubated in blocking solution (5% donkey serum, 0.25% triton, and 0.05% sodium azide in 0.01 M PBS) for 1 hour before left to incubate in primary antibody for 72 hrs. Then, sections were rinsed six times in 1x PBS and then allowed to incubate in an anti-rabbit GFP secondary antibody (anti-rabbit-GFP, Life Technologies, 1:800) for four hours. Final rinses and Dapi (ThermoFisher Scientific, 1:1000) staining were achieved before tissue was mounted on glass slides and cover-slipped.

### Quantitative image analysis

The Prelimbic Cortex (PL), Infralimbic Cortex (IL), Dorsal Striatum (DS), Nucleus Accumbens (NAc), Paraventricular Nucleus of the Thalamus (PVT), Central Amygdala (CeA) and Basolateral Amygdala (BLA) were imaged with a Zeiss LSM 780 confocal microscope equipped with a 63x oil lens (NA 1.4). Neuroanatomical landmarks were visualized by transmitted light such as the external capsule, which was used as the lateral border to guide the experimenter to the amygdala region. Representative sampled regions are delineated in **Figure 2** (A-F) with panels D, and F showing robust Fos-expressing cortical, amygdalar, and paraventricular thalamic activated neurons. A tiling method with a 10% overlap between adjacent images was employed to encompass the entire brain region of interest in the final tiled stack image. Targeted brain regions were imaged with four image stacks acquired for each brain region sampling across the rostrocaudal axis. Each image stack was acquired with a z-step size of 0.5µm and post-processed to generate a maximum intensity projection (MIP). Regions of interest were contoured using the freehand contouring method in NeuroInfo (MBF Bioscience). For systematic analysis, the same contour with the same surface area dimensions was applied for each respective brain region in subsequent MIP images.

**Figure 2.**
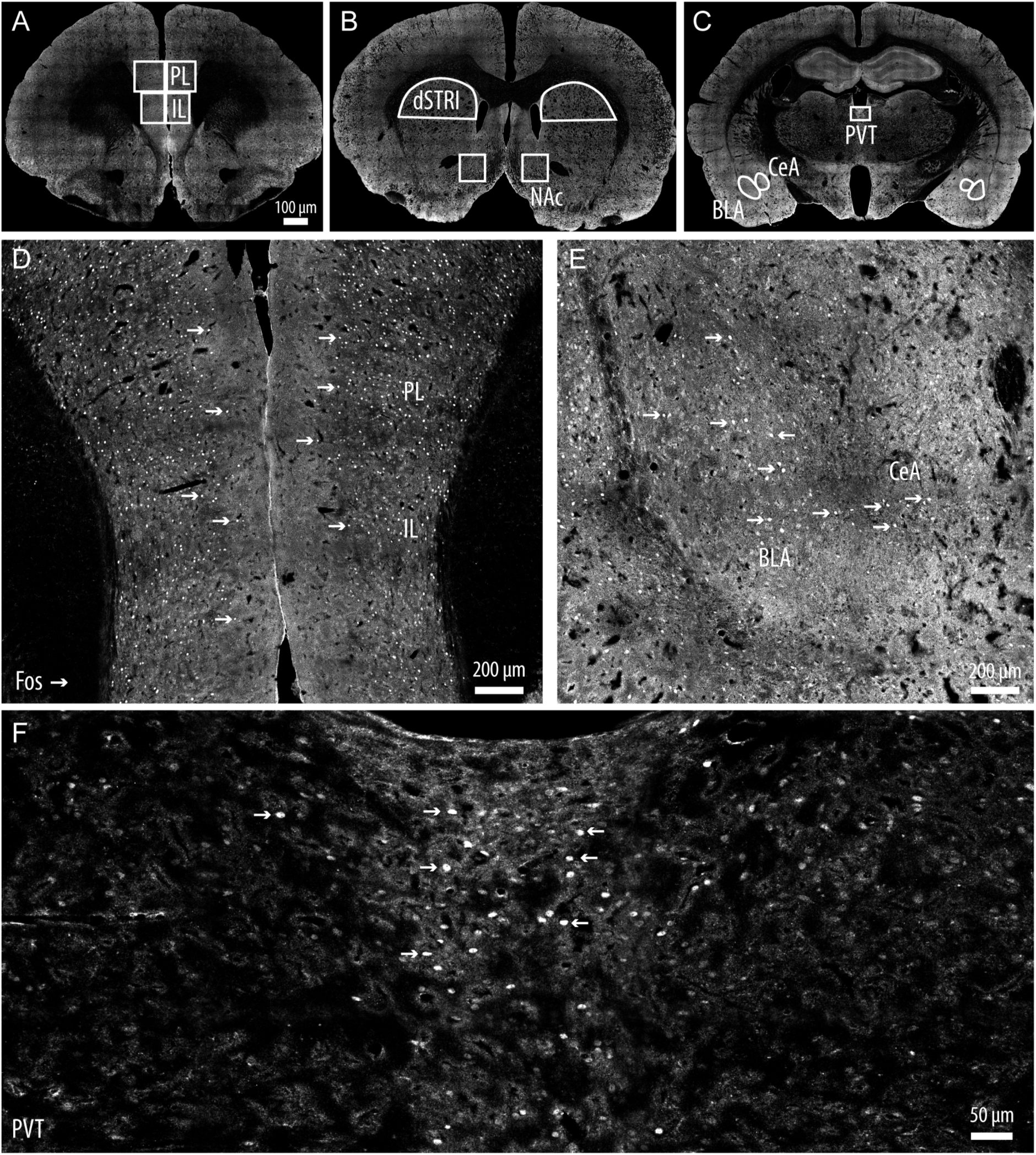
Representative confocal images depicting brain regions sampled for Fos-positive quantitative analysis of stimulus-activated neurons within engrams. **(A)** Prelimbic cortex (PL) and Infralimbic cortex (IL). **(B)** Dorsal Striatum (DS) and Nucleus Accumbens (NAc). **(C)** Paraventricular Nucleus of the Thalamus (PVT), Central Amygdala (CeA) and Basolateral Amygdala (BLA). **(D)** Arrows denote representative Fos-positive of the PL, IL, CeA, BLA in **(E)** and PVT active neuronal ensembles in **(F)**.

Fos-expressing neurons were detected automatically with the cell detection function in NeuroInfo. First, a diameter between 7 – 20 µm was set in order to detect these activated neurons with an average diameter of 9 µm. The stringency of the automatically detected cells was then adjusted in the software in order to reduce false positives as well as to detect neurons with lower Fos-expression. Fos labeling has a large dynamic range with robustly labeled neurons as well as more faintly labeled neurons. In order to detect low expressing Fos-positive (Fos+) cells and to remove artifacts, AI cell detection was employed. For this, an AI-driven neural network was trained using training data from approximately 200 regions selected randomly from a pool of 40 whole brain images. This training data was generated by humans confirming, removing, or adding markers on presumed Fos-positive neurons detected via a standard parameterized cell detection algorithm (a modified LOG detector) configured to detect as many cells as possible (i.e., all true-positive cells and some false-positive cells were both detected and then the results were curated by a human observer). This semi-automated method enabled the labeling of over 20,000 cells. This training data was used to augment our existing classifier to better recognize the broad expression of Fos protein while also increasing its ability to reject artifacts, such as the edges of vessel, section edges, and ventricle intersects (**Figure 3**).

**Figure 3.**
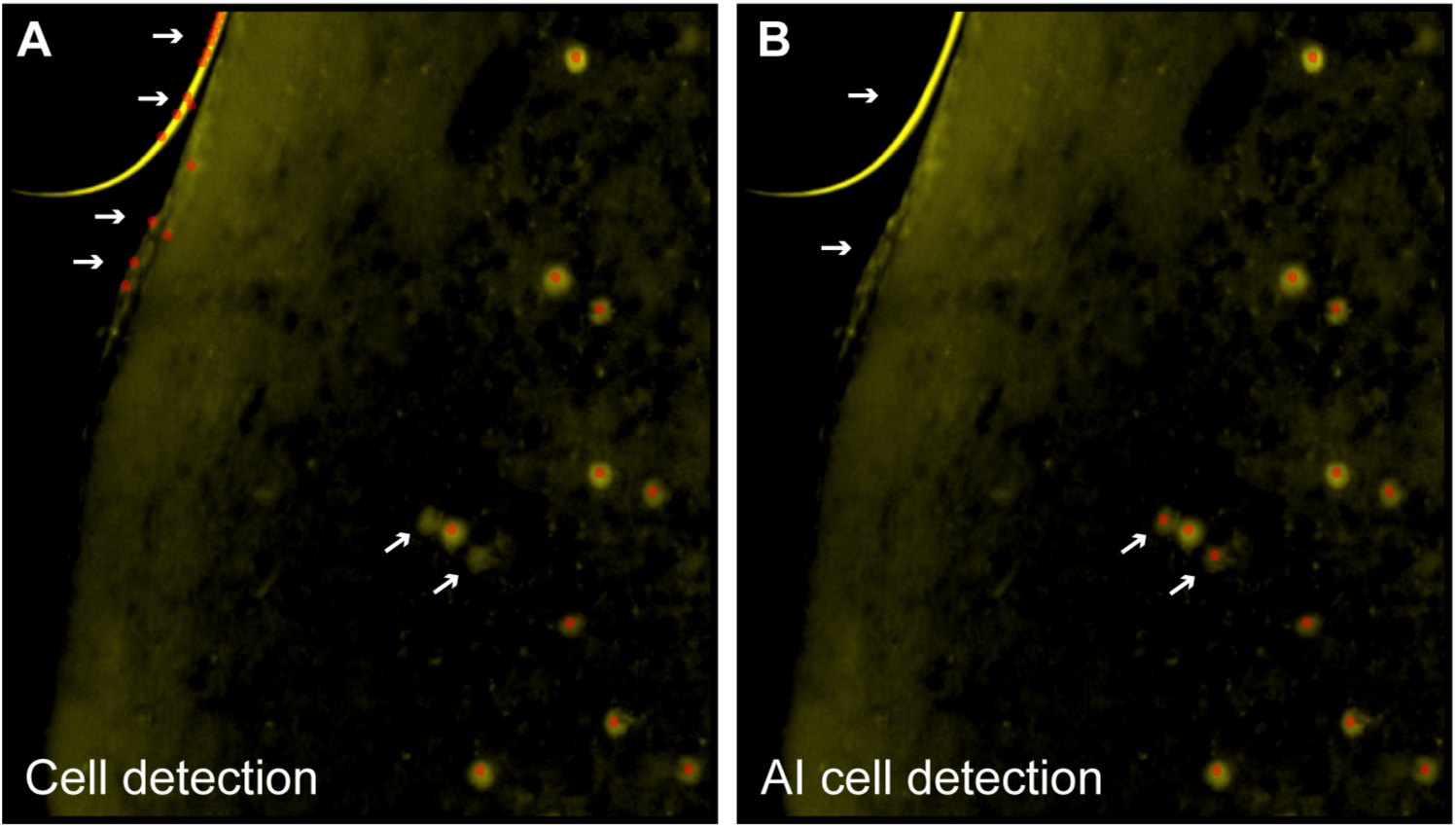
Images showing Fos activated neurons using a standard cell detection methods **(A)** vs an improved cell detection method using Artificial Intelligence with Machine Learning to eliminate false positives due to edge effects and capture dimmer active neurons **(B)**.

All automated cell detections were validated and edited by an experimenter before finalization. Three independent experimenters edited the automated cell detections and all three finalized cell count values were averaged. Experimenters were blind to the treatment condition. Fos cell counts were normalized over the total surface area of each region of interest to obtain Fos-density (Fos counts/mm^2^) as a proxy for neuronal activation representing engrams (Josselyn and Tonegawa 2020; Curran and Morgan 1985; Greenberg and Ziff 1984; Guzowski et al. 2001).

### Statistical analysis

The design consisted of four experimental groups: (1) DEP-WDL, (2) NDEP-WDL, (3) DEP-NWDL, and (4) NDEP-NWDL as described above. To study the effects of the respective contextual stimulus exposure on neuronal activity within engrams, Fos density was used as the dependent variable. While the assumptions underlying parametric tests were met for independent random sampling and homogeneity of variance, normal distributions were not observed in these data (Cohen 2008). For this reason, the treatment effects on Fos density were analyzed by non-parametric tests and where appropriate followed by post hoc comparisons with Bonferroni corrections.

## RESULTS

### Overall neuronal activation within engrams is increased in DEP-WDL relative to non-depended groups

To examine whether WDL stimulus exposure had an effect on neuronal activity within engrams, Fos density was quantified in key brain regions with an established role in alcohol seeking and craving. A Kruskal-Wallis test was conducted to examine the differences in Fos density among the following four groups: DEP-WDL, NDEP-WDL, DEP-NWDL, and NDEP-NWDL. The results of this test indicated a significant difference in Fos density among the treatment groups (H (3) = 7.5; P < 0.05). Post-hoc analysis by pairwise comparisons revealed an increased overall density of activated neurons in the DEP-WDL group compared to both non-dependent groups (**Figure 4A**, DEP-WDL, Dunn’s test with Bonferroni correction P < 0.05). No significant difference was found between the DEP-WDL and DEP-NWDL (n.s.) or between DEP-NWDL and the non-dependent groups (n.s.).

**Figure 4.**
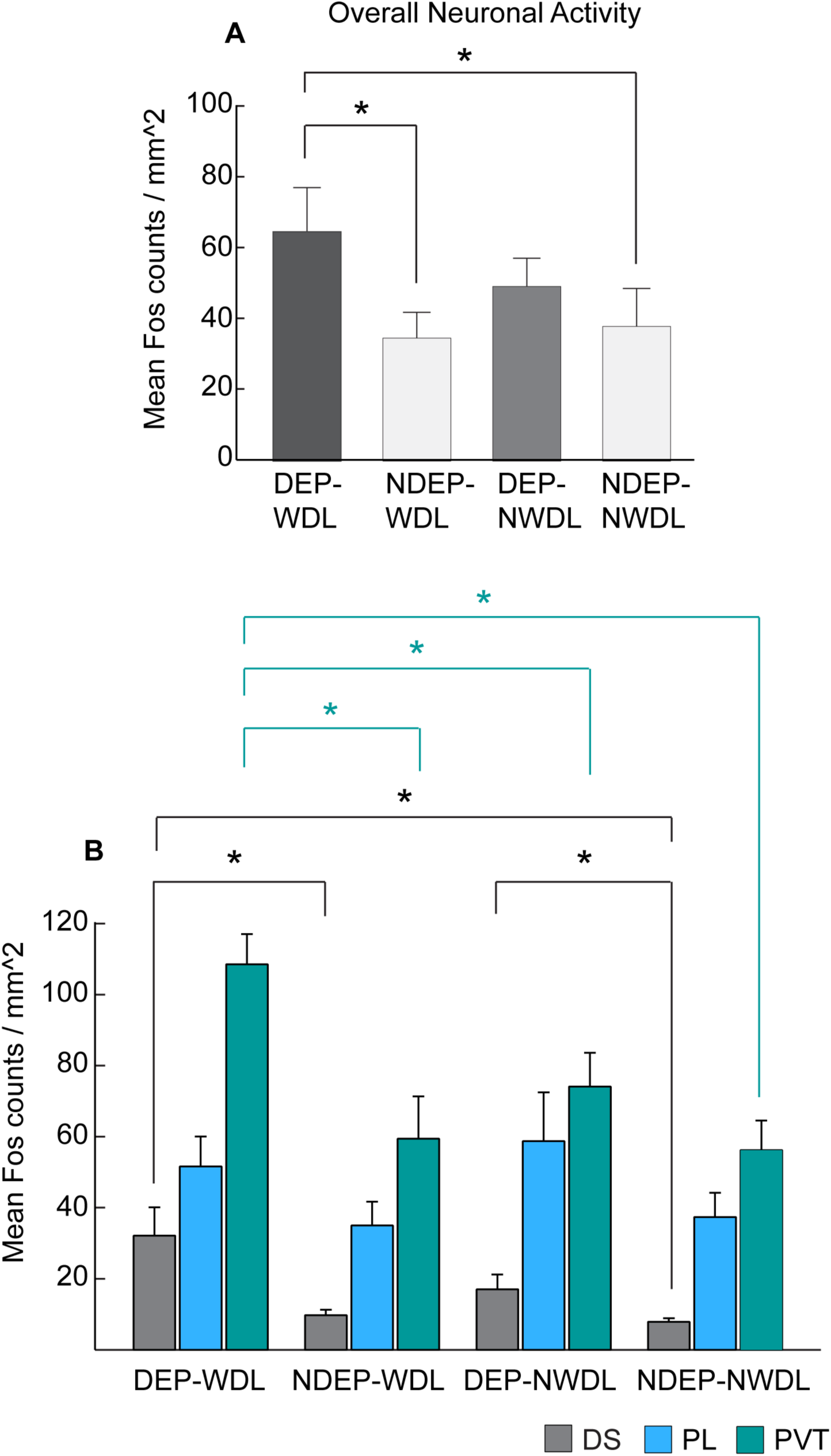
**(A)** Plot summarizing the overall Fos-activated neurons within engrams for all four experimental groups DEP-WDL (N = 13), NDEP-WDL (N = 13), NDEP-POST (N = 9) and DEP-POST (N = 9). **(B)** Measurement of Fos activity in the DS (n = 15), PL (n = 14) and PVT (n = 15) revealed an increased number of activated neurons within engrams in the PVT of DEP-WDL rats (Kruskal-Wallis, Dunn’s test with Bonferroni correction P < 0.05). Increased neuronal activity representing a DS engram was also present in both dependent groups (DEP-WDL, DEP-NWDL) relative to the non-dependent (NDEP-WDL, NDEP-NWDL) groups (Kruskal-Wallis, Dunn’s test with Bonferroni correction P < 0.05. Bars represent mean and SEM).

### Learning experience-dependent neuronal activation within engrams is increased in PVT and DS

To better understand the neuroanatomical localization of engrams reflected by neuronal activation changes, Fos density was initially analyzed in three neuroanatomical brain regions with an established role in alcohol/drug addiction: dorsal striatum (DS), prelimbic cortex (PL), and paraventricular nucleus of the thalamus (PVT) (Zhou and Zhu 2019; Belin and Everitt 2008; Nedelescu et al. 2022). A Kruskal-Wallis test was conducted to examine the differences in neuronal activation among the four treatment groups and across the three brain regions. The results revealed a significant treatment group difference in Fos density in the DS and PVT (DS: H (3) = 8.5, P < 0.05; PVT: H (3) = 10.5, P < 0.05) but not in the PL (H (3) = 4.4; n.s.). Multiple pairwise comparisons by Dunn’s test with Bonferroni correction revealed that the DEP-WDL group showed an increased density of activated neurons specifically in the PVT compared to all other treatment groups (**Figure 4B**, asterisks denote P < 0.05). In addition, increased neuronal activation was observed in the DS of DEP-WDL rats relative to the non-dependent groups, NDEP-WDL and NDEP-NWDL (P < 0.05); however, no difference was found between the DEP-WDL and DEP-NWDL for this striatal region (**Figure 4B**). Since these data revealed that WDL experience was a critical factor in the observed engrams represented by neuronal activity increases, specifically in PVT, we focused the remaining analyses on activity-dependent neuronal changes between the DEP-WDL and NDEP-WDL groups.

For a more comprehensive analysis across different neuroanatomical brain regions, Fos counts were extended to four additional brain regions with an established role in drug addiction including the IL, NAc, CeA, and BLA (**Figure 5A**). To gain further insights about the functional changes across neuroanatomical regions and recruited engrams as a result of the respective learning experience during the withdrawal state in dependent animals vs. non-dependent controls, a Mann-Whitney U test was used to compare neuronal activity in DEP-WDL and NDEP-WDL across all eight brain regions: DS, PL, PVT, IL, NAc, CeA, BLA, and LH. The results revealed a significant increase in the density of activated neurons between DEP-WDL and NDEP-WDL following WDL experience in the PVT, DS, and CeA (PVT: z = 1, P < 0.05; DS: z = 0, P < 0.05; CeA: z = 0, P < 0.05), but no difference in Fos density for the PL, IL, NAc, BLA or LH between the DEP-WDL and NDEP-WDL groups (**Figure 5A**).

**Figure 5.**
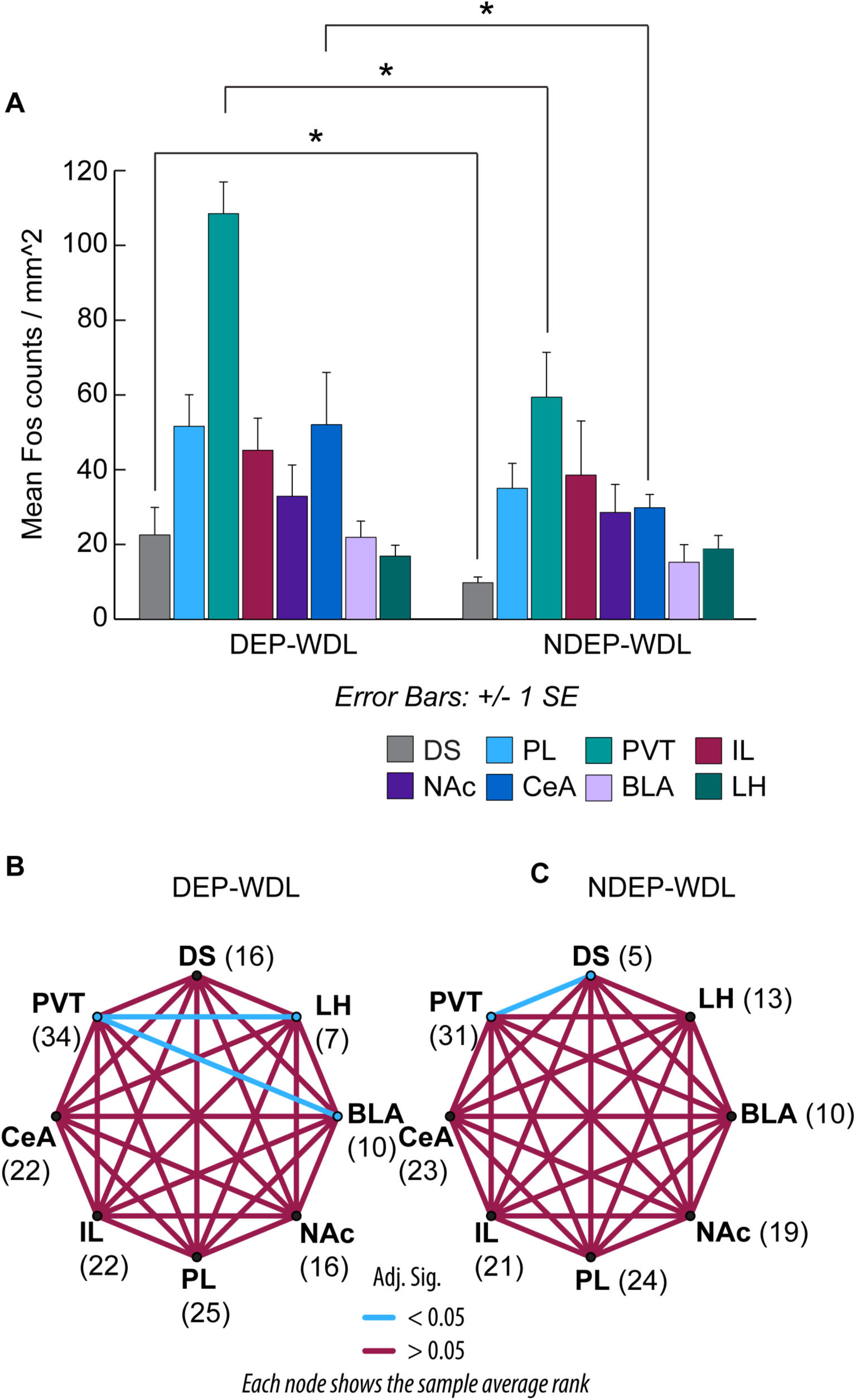
**(A)** Recruitment of engrams in the DS, PVT and CeA, but not in the IL, PL, NAc, BLA or LH following WDL (Mann-Whitney U, DS, PVT, CeA P < 0.05). **(B, C)** Pairwise comparisons confirmed deferentially recruited engrams among brain regions in the DEP-WDL vs. the NDEP-WDL conditions. Nodes represent the different brain regions analyzed for Fos protein expression. The number adjacent to each node represents the mean rank of the Kruskal-Wallis test. The edges connecting the nodes represent the pairwise comparisons between the groups. Blue lines show statistically significant differences between brain regions. Comparisons in these plots evaluate whether differences in ranks are statistically meaningful: higher ranks (e.g., PVT 34.0) indicate a higher average Fos density compared to lower ranks (e.g., LH with a rank of 7.0). Regions with similar ranks are connect by red lines. Regions with highly disparate ranks are connected by blue lines. **(B)** In the DEP-WDL group, a PVT engram is observed by the significantly increased Fos activation compared to both the BLA and LH, but not compared to the DS. **(C)** In the NDEP-WDL group, a PVT engram is observed by significantly increased Fos activation in the PVT compared to the DS.

**Figure 6.**
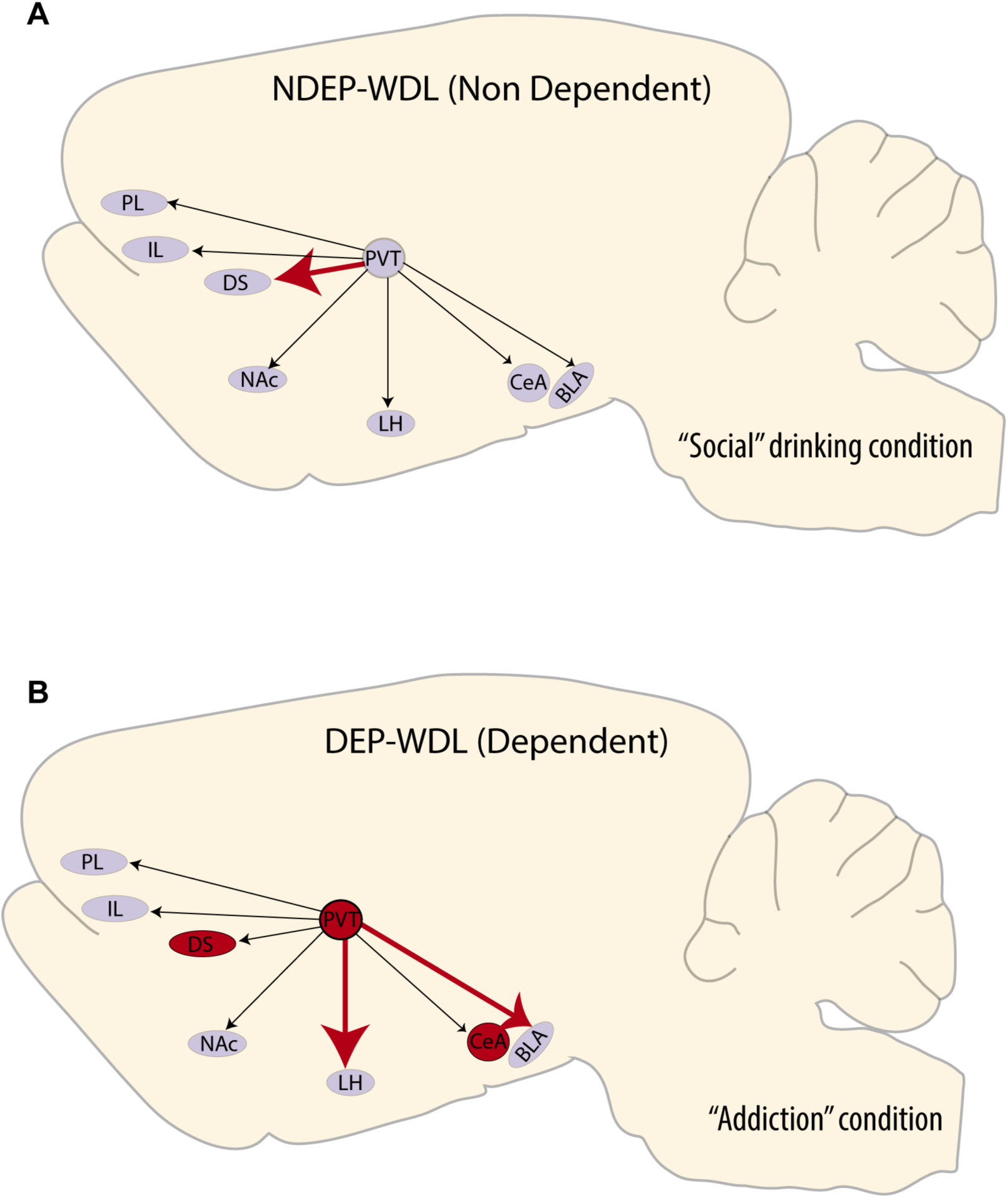
Illustration of the formation of engrams associated with context-induced alcohol seeking in rats with WDL experience and “dependent” drinking **(B)** vs alcohol seeking induced by the same context in nondependent rats with a “social drinking” history **(A)**. Arrows represent PVT neurons projecting to target brain regions analyzed in this study. Thick red arrows represent within group neuronal activation pattern changes as a consequence of developing dependence and allostasis (NDEP-WDL vs DEP-WDL). Red regions represent the formation of engrams between the “addiction” condition (DEP-WDL) and “social drinking” condition (NDEP-WDL). These differential changes across the different neuroanatomical regions reflect a functional reorganization within the brains of animals with a WDL (DEP-WDL) history.

### Within group comparisons

Engrams analyzed by pairwise comparisons within the NDEP-WDL group showed that neuronal activity was significantly increased in the PVT compared to the DS, but not statistically different compared to the remaining brain regions analyzed including the LH, BLA, NAc, PL, IL or CeA (Pairwise comparisons with Bonferoni correction, **Figure 5C**, P < 0.05). By contrast, analysis of engrams in the brains of DEP-WDL animals showed that neuronal activation was significantly higher in the PVT compared to both the LH and BLA but statistically not significantly different compared to the DS, NAc, PL, IL, and CeA (**Figure 5B**). Comparisons in these plots evaluate whether differences in ranks were statistically significant [(blue lines = statistically significant; red lines = n.s.). Higher ranks (e.g., PVT: 34.0) represent a higher average Fos density than lower ranks (e.g., LH: 7.0), and thus, suggestive of the recruitment of an engram.

## Discussion

### Differential recruitment of engrams in alcohol seeking subjects with WDL experience

The findings confirm a significant differential pattern of engrams recruited during context-induced compulsive alcohol seeking in rats with a WDL history than in rats without this history and that do not show compulsive behavior (Kozanian et al. 2022). More specifically, engrams encoding associations between environmental stimuli linked to the reversal of adverse withdrawal effects (i.e., negative hedonic states) were identified in the DS, PVT, and CeA (**Figure 4B**, **5A**). In contrast, exposure to stimuli conditioned to the hedonically positive aspects of alcohol consumption in nondependent or postdependent rats without WDL experience did not activate engrams in the PVT and CeA regions and produced only mild, albeit significant, activation in the DS in postdependent rats. These observations support the hypothesis that the conditioned effects of contextual stimuli associated with the reversal of withdrawal distress are differentially represented in the brain compared to the effects of stimuli associated with all other learning conditions in both alcohol dependent and nondependent rats.

The prominent engram identified exclusively in the PVT of the WDL group suggests that the PVT plays a major role in the acquisition of the negative contingency between alcohol consumption and the dysphoric effects of withdrawal and the resulting development of compulsive drug seeking (Kozanian et al. 2022). The PVT is a key hub of neural circuits implicated in drug addiction (Dayas et al. 2008; Matzeu et al. 2017; Zhou and Zhu 2019). Moreover, this nucleus has an established role in emotional responses to anxiety and stress (Hsu et al. 2014; Tang et al. 2024). Stress increases neuronal activity in the PVT (Fiedler et al. 2021) and the PVT was shown to be critical for stress-induced reinstatement of oxycodone seeking (Illenberger et al. 2024). Dysphoria as in alcohol withdrawal is strongly associated with stress (Koob and Le Moal 2001; Merlo Pich et al. 1995). Therefore, the results may point to a major role of stress in WDL learning and compulsive drug seeking following WDL acquisition. The stress-sensitive PVT area projects axonal fibers to the CeA, a region associated with negative emotion, stress, alcohol dependence, and particularly the aversive effects of alcohol withdrawal (Roberto et al. 2021; Merlo Pich et al. 1995). Exposure to the WDL-associated stimulus context resulted in the recruitment of a CeA engram, suggestive of increased CeA neuronal activity compared to CeA neurons in NDEP-WDL subjects in which stimulus context was associated only with the hedonically positive experience of alcohol consumption (**Figure 5A**) (Kozanian et al. 2022). Chemogenetic inhibition of the PVT to CeA projection alleviates stress responses (Zhao et al. 2022), supporting a role of this projection system in stress.

### Dorsal striatum engrams and dependence

The development of dependence is associated with the emergence of habitual behavior which is thought to be mediated by striatal brain regions, in particular the DS (Barker et al. 2015; Everitt and Robbins 2013; O’Tousa and Grahame 2014; Belin and Everitt 2008). Thus, the observed DS engram in the WDL group is possibly explained by a progressive engagement of dorsal striatal regions in habit formation that emerges during the development of substance dependence as proposed by the “spiraling” hypothesis [(i.e., processes through which the ventral the ventral striatum comes to exert control over dorsal striatal processes mediated by so-called “spiraling,” striato-nigro-striatal, circuitry (Haber et al. 2000; Ikemoto 2007; Belin and Everitt 2008). A DS engram was also recruited in the brains of postdependent (DEP-NWDL) animals (**Figure 4B**, **5A**). These rats had a history of dependence but without WDL experience and were trained to associate the stimulus context with alcohol availability following completion of alcohol withdrawal (Kozanian et al. 2022). The DS engram activation in these animals was smaller than in the WDL history group but significantly different from nondependent controls (NDEP-NWDL). Thus, DS engrams are encoded and become activated to some degree not only as a consequence of the WDL experience or negative reinforcement learning, but also other dependence associated or experiential factors independent of WDL.

### Intragroup engrams

The within-group pairwise comparisons across the eight brain regions within the DEP-WDL versus NDEP-WDL groups revealed differential patterns of engrams across brain regions for each condition. A recruited engram was found within the NDEP-WDL control group in the PVT region which showed increased neuronal activation relative to the DS region (**Figure 5C**). However, the PVT engram within the experimental DEP-WDL group differed in activation relative to both the LH and BLA, suggestive of a re-organization in inter-region activity patterns associated with WDL experience (**Figure 5B**). That the PVT and DS were not differentially activated in the DEP-WDL group is the result of increased neuronal activity and emergence of the DS engram within these subjects (i.e., DS, with rank 16 in the DEP-WDL condition vs DS with rank 5 in the NDEP-WDL group). This change within the brains of DEP-WDL animals is possibly explained by the engagement not only of stress regulatory systems (in the PVT and CEA) but also of dorsal striatal regions implicated in habit formation that emerges during the development of substance dependence as proposed by the “spiraling” hypothesis discussed above (Haber et al. 2000; Ikemoto 2007; Belin and Everitt 2008).

### Addiction as a state of reward dysregulation

Non-dependent (NDEP-WDL) rats experienced a positive hedonic state during alcohol self-administration. In contrast, dependent (DEP-WDL) rats experienced a profound negative hedonic state during withdrawal. According to the opponent-process hypothesis of motivation an initial positive hedonic A process gives rise to an opposing negative hedonic state or B process below a baseline state or hedonic equilibrium (Solomon and Corbit 1974; Solomon and Corbit 1978). This opponent-process is strengthened through use and weakened through disuse. Multiple activation of the initial A process, such as by repetitive use of alcohol, leads to a reduction of this positive hedonic A process and an opposing strengthening of the B process below baseline equilibrium. According to this hypothesis, the B process represents an adaptive response by which the neural system minimizes deviations from hedonic equilibrium. Over time, continued use of alcohol (or other hedonically positive stimuli or substances) leads to an allostatic state in which the A and B processes still occur but with both falling below the original equilibrium into negative hedonic territory (Koob and Le Moal 2001; Koob et al. 1989).

With respect to the present findings, the reversal of the dysphoric withdrawal states by alcohol consumption in the DEP-WDL group presumably conveyed potent incentive value to the drug, driving further alcohol consumption and exacerbating the negative hedonic B process such that the drug further gained in hedonic valence over the progression toward the allostatic state. This progressive deviation from hedonic equilibrium and the transition to a pathological allostatic state is known to be associated with corresponding adaptive changes in brain reward systems (Breese et al. 2005; Zorrilla et al. 2012; Koob et al. 1994; Koob et al. 1998; Weiss et al. 1996; Patel et al. 2022). Similarly, associative processes that link environmental contexts with the withdrawal-reversing effects of alcohol are likely to progress toward the engagement of different neuronal systems compared to those recruited when alcohol acts as a positive reinforcer. The PVT and CeA engrams were associated specifically with WDL experience and likely directly linked to the reversal of the stressful aspect of alcohol withdrawal in rats with WDL experience. By contrast, the DS engram, albeit differential, was observed both in dependent animals with WDL experience (DEP-WDL) and animals with a history of dependence (DEP-NWDL). The engrams identified here may serve as a neural substrate for compulsive drug seeking resulting from (1) negative reinforcement learning associated with hedonic allostasis (PVT), (2) the development of habitual behavior over the progression of dependence (DS), and (3) stress memories associated with the stimulus context in which withdrawal and reversal of withdrawal was experienced (CeA). To confirm a role for these engrams specifically in mediating WDL experience, future experiments involving silencing of each engram with cell-type specific methods in the DEP-NWDL and NDEP-NWDL groups would be required (Cruz et al. 2013). Additionally, these findings may have broader implications for maladaptive behavior linked to reward dysregulation beyond substance use disorders. Research to establish the neurobiological alterations that link the conditioned incentive value of substance-associated contexts and their role in exacerbating drug seeking could be extended in the future to other types of maladaptive behavior, more generally. These include, but are not limited to, systems that regulate fear-conditioning, anxiety disorders (Nedelescu et al. 2010; Hagihara and Luthi 2024; Moscovitch and LoLordo 1968), traumatic avoidance learning (Solomon et al. 1953) and possibly predatory behavior.

### Study Considerations

Neuronal activation to identify engrams was measured by Fos protein expression, which has been utilized as a marker for recent neuronal activity (Pettit et al. 2022; Cruz et al. 2013; Cruz et al. 2015). Neuronal ensembles are sparse patterns of neurons that are selectively activated by environmental stimuli, whereas engrams represent the long-lasting cellular alterations that physically encode learned associations within neuronal ensembles (Cruz et al. 2015; Cruz et al. 2013; Josselyn and Tonegawa 2020; Eichenbaum 2016). The present study was designed to identify engrams that encode withdrawal-related learning (WDL) in dependent animals and subsequently drive substance seeking and craving. Because the WDL procedure requires the conditioning of contextual stimuli associated to alcohol availability and the subjective effects of alcohol consumption over multiple exposures for three weeks, it is reasonable to assume that the neuronal activation induced by re-exposure to these stimuli reflects the long-lasting neuronal changes that represent an organizational neural response or engrams.

## Author Contributions

H.N. conceived the project, performed the histology and imaging with assistance from E.M., N.R., A.G., and R.B. Fos+ neurons were counted by E.M., N.R., and A.G. using a protocol designed by H.N. and N.O. and supported by MBF Bioscience. H.N. conducted the formal data analyses and wrote the manuscript with critical input from F.W. Microscopy costs were covered by N.S. Finally, F.W. secured the funding and supervised the project with H.N. as lead project administrator and supervisor.

## Acknowledgments

We thank Olga Kozanian for conducting the behavioral experiments and for ongoing dialogue during the preparation of this manuscript. This work was supported by a K01 DA054449 (H.N.), R01 AA023183 (N.S.), U01DA055017 (N.S.), R01 AA027555 (F.W.).

